# Extraction, purification and anti-TMV effects of α (β)-2,7,11-cembratriene-4,6-diol from tobacco leaves

**DOI:** 10.1101/2021.06.01.446537

**Authors:** Yongxin Feng, Haijiang Jia, Hui Guan, Weidong Zhang, Yanbin Zhou, Kaiyi Liu, Yuejing Wang, Qiuli Li, Weihua Chen, Muhammad Aamir Sohail, Jie Wang, Shen Huang, Jianyu Wei

**Author notes:** Correspondence (S.-H); (J.-Y.W.). Both authors made equal contributions to this work (Yongxin Feng, Haijiang Jia).

## Abstract

**Background:** Acetone ethanol extracts from tobacco leaves have antiviral activity against TMV, but the antiviral effects of their specialized metabolites have not been systematically studied yet, especially the underlying mechanism is still unexplored.

**Results:** The tobacco cembranoids α(β)-2,7,11-cembratriene-4,6-diol (α(β)-CBD) were extracted and purified with an effective and green protocol including innovatively added 5% phosphoric acid for elution, one time silica gel chromatographic column separation and impurity removal and further HPLC purification. The results of antiviral activities against tobacco mosaic virus (TMV) with the local lesion counting method showed that α(β)-CBD have *in vivo* higher protective effects of 73.2% and 71.6%, at 75.0 μM, respectively, than control agent Ningnanmycin (53.1%). Notably, The results of ELISA and and TMV-GFP fluorescent optical imaging assay indicated a obviously reduced viral protein and weaker GFP fluorescence signal and smaller infection area, which confirmed their anti-TMV activities at protein level. Furthermore, the enhanced production of SA and JA and the significantly increased transcription of of JA signaling pathway (*COI1* and *PDF1.2*) and SA signaling pathway genes (*PR1*, *NPR1* and *EDS1*) in treated plants further conformed that exogenously applied α(β)-CBD can effectively elicit the tobacco plant immunity against TMV.

**Conclusions:** The α(β)-CBD mainly stimulates disease resistance of tobacco plants to resist TMV and it can be used as bioagents to control TMV in the future.

## 1. Introduction

*Nicotiana tabacum* L. (the flue-cured tobacco), an herbaceous plant belonging to the *Nicotiana* genus (Solanaceae family), is now cultivated worldwide and has long been used medicinally and recreationally as the most economically important industrial crops[1,2]. Moreover, *Nicotiana* species are commonly investigated for the biological activity of their specialized metabolites in responses to abiotic stress or biotic stress factors such as pathogens[3,4]. Tobacco mosaic virus (TMV) is the most ancient virus that causes massive economic losses to tobacco, pepper, cucumber and ornamental crops globally. Several findings suggested that acetone ethanol extracts from *N. tabacum* leaves have antiviral activity against TMV[5–8], but the antiviral effects of their specialized metabolites have not been systematically studied yet, especially the underlying mechanism is still unexplored.

Cembranoids, a group of natural diterpenoid products comprising four isoprene units of a 14-carbon macrocyclic skeleton, are mainly distributed in *Nicotiana tabacum*, *Pinus genera*, and some marine species (e.g., soft coral)[9,10]. In *Nicotiana* spp., cembranoids are most prevalent in the surface secretion of leaves and flowers[11]. A wide variety of bioactivities, including neuro-protective, anti-cancer, anti-invasive, antifungal, antiviral and anti-Inflammatory activities against animal viruses, have been carried out with two main cembranoids, α-cembratrien-diol (α-CBD) and C4 epimer (β-cembratrien-diol, β-CBD) (Figure 1)[12–14]. In addition to exhibiting suppression in spore germination of many fungal diseases (*Peronospora tabacina, Colletotrichum lagenarium, Candida albicans, Fusarium chlamydosporum, Aspergillus niger and Alternaria alternata*), exogenously applied α(β)-CBD also inhibited the growth of bacteria (*Staphylococcus aureus*, *Bacillus subtilis* and *Proteus vulgaris*)[15,16]. Interestingly, in response to tobacco mosaic virus infection, a substantial increase in cembranoids, in particular α/β-CBD, has been achieved with systemic acquired resistance (SAR) leaves[17,18]. However, it is scarcely examined whether the exogenous application of α(β)-CBD has inhibitory effects on tobacco virus infection or not.

**Figure 1.**
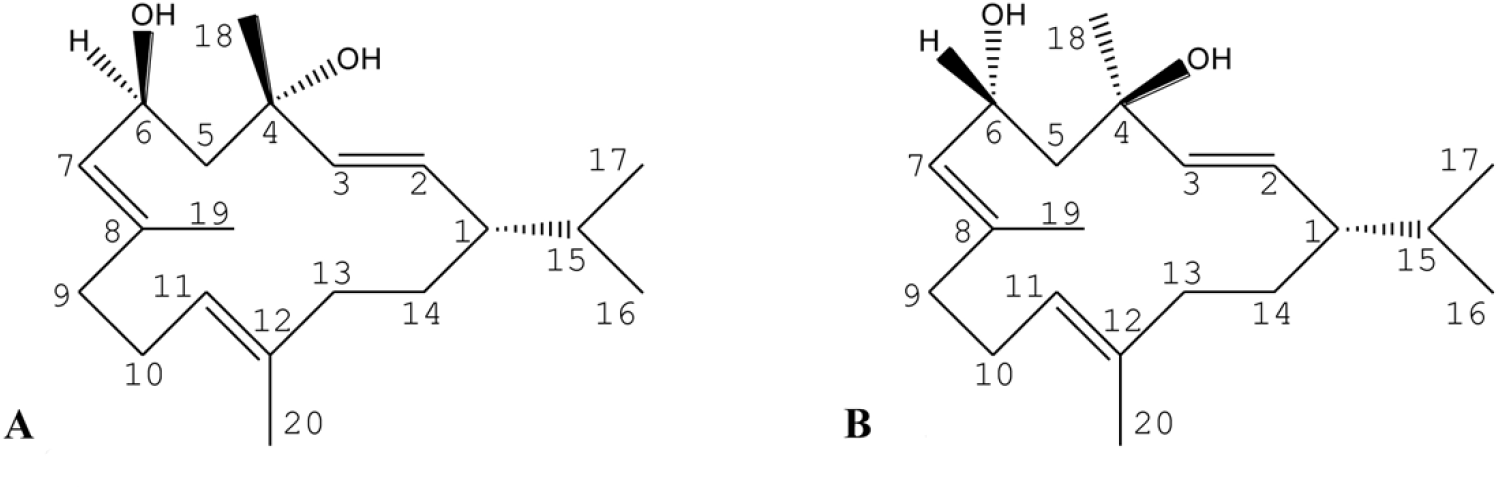
Molecular structure of α-CBD (A) and β-CBD (B).

In continuing efforts to identify natural product-derived antiviral agents against plant virus, we herein report the simultaneous extraction and identification of α-CBD and β-CBD from Yunyan 100 (a variety of *N. tabacum*). TMV-GFP based systemic host approach was used to assess the two compounds’ anti-TMV potential in tobacco. The isolation, structural identification and biological evaluation of these compounds are listed here.

## 2. Materials and methods

### 2.1. Plant material

Fresh mature leaves of tobacco (Yuyan 100) for purification of α-CBD and β-CBD were harvested in Xuchang District (33.53 north latitude, 113.49 east longitude), Henan Province, China, in July 2014. Tobacco plants were cultivated in a light chamber at the light intensity of 10,000 lux with a long day photoperiod (18 h : 6 h, light : dark). The following experiments were conducted when *Nicotiana tabacum* L. cv. Samsun *NN* was selected at the seventh leaf stage for the local lesion counts, and *N.benthamiana* had grown 6 leaves for systemic TMV infection determination. TMV-GFP (a recombinant TMV encoding green fluorescent protein) was a gift from Professor Yule Liu, (Tsinghua University, Beijing, P. R. China).

### 2.2. Large scale purification and isolation

The purification and isolation of α(β)-CBD was conducted as described previously with some revision[19,20]. The leave extracts from fresh mature tobacco leaves (200 kg) were extracted with dichloromethane (CH_2_Cl_2_) three times (3-5 sec each time at room temperature). The combined extracts were filtered to remove insoluble ingredients and green transparent filtrates were obtained. A brown residue (0.8 kg) was obtained after the removal of the solvent under reduced pressure at 45 °C, and completely dissolved under ultrasound radiation conditions in CH_2_Cl_2_. CH_2_Cl_2_ extract was applied to silica gel (100-200 mesh) column chromatography eluted with MeOH : C_4_H_8_O_2_ =3 : 1, and the solvent was distilled under reduced pressure, and1 resuspended with CH_2_Cl_2_ (0.4 kg). CH_2_Cl_2_ dissolved matter was passed through two silica gel column chromatography of 200-300 mesh and 300-400 mesh in order eluted with CH_2_Cl_2_ : C4H_8_O_2_ = 3 : 1. α-CBD (0.07 kg) with 70% purity and β-CBD (0.07 kg) with 70% purity were obtained. Finally, α(β)-CBD (0.07 kg) with 70% purity were applied C18 column chromatography (YMC-Pack ODS-A 250 × 20 mml.D.S-5um) eluted with C_2_H_3_N : H_2_O =3 : 1. And α(β)-CBD with 98 % purity were obtained.

The UPLC conditions for detecting CBD after purification were: capillary column: ACQUITY UPLC BEH C18 column (Waters, Milford, USA), 1.7 μm (2.1 × 50 mm); injection volume was 1 μL; the mobile phase was a mixture of acetonitrile and water in a ratio of (60:40) at a flow rate of 0.5 mL/min; column temperature was 40 °C, and the UV wavelength was set at 210 nm.

Purified α(β)-CBD was checked by GC-MS (Agilent 7895A-5975C, USA). Chromatographic conditions were: capillary column: DB-5 (60 m × 250 μm × 0.25 μm); carrier gas: He; injection volume: 1 μL; injector temperature: 180 °C; split ratio: 6:1; column temperature 120 °C for 1 min, programmed at 5 °C/min to 180 °C, 2 °C/min to 240 °C, and 5 °C/min to 280 °C. Mass spectrometer conditions were: interface temperature: 280 °C; quadrupole temperature: 150 °C; ion source temperature: 250 °C; solvent delay 8.5 min; mass scan range: 35–500 AMU.

After purification of the C18 column, we obtained two peaks containing CBD. The colorless compound 1 from the first peak was condensed to dryness and dissolved at 55 °C in N-pentane. Compound 1 solution was held at 25 °C, and was slowly transformed into the crystals. Compound 2 from the second peak was white and concentrated to dryness and dissolved in N-pentane at 55 °C. Compound 2 solution was kept at −5 °C, and was slowly transformed into the crystals. High-resolution GC-MS for determining α(β)-CBD is described in 2.3. NMR for resolving the structure of α(β)-CBD was done[21]. ^1^H and ^13^C experiments were carried out at 600 MHz on a Brucker Advance 600 MHz spectrometer (Brucker, Rheinstetten, Germany)[22].

### 2.3. Antiviral activity assays

TMV was According to Gooding’s method[23], TMV virions maintained in *N. tabacum* cv. K326 were extracted and purified from the harvested leaves of systemically inoculated seedlings by homogenization in 0.01 M PBS (phosphate-buffered saline) buffer. The supernatants obtained by centrifugation at 1800 *g* for 3 min were stored at – 20 °C for use.

Purified α(β)-CBD was dissolved in DMSO and diluted with distilled H_2_O containing 0.1% Tween 80 to the required concentrations (12.5, 25.0, 50, 75 μg/mL) for the following treatments. Ningnanmycin (200 μg/mL) was used as a positive control.

For the protective assay *in vivo*[24,25], The right half leaves of *N. tabacum* L. cv. Samsun *NN* were gently rubbed with α and β-CBD solution, respectively. After 6 h, the same area received the inoculation of 100 μL TMV (10.19 μg/mL), The positive control was the left half leaves smeared with TMV after applying DMSO solvent. The numbers of local lesions were counted 5 days after inoculation. Five replicates were implemented for each sample.

For the inactive assay *in vivo*[24,25], the TMV particles (10.19 μg/mL final concentration) were firstly mixed with the tested agents solutions at room temperature for 15 min, and then the mixed solution was spread on the left half leaves of *N. tabacum* L. cv. Samsun *NN*, whereas the right side was smeared with TMV mixed with DMSO solution. The numbers of local lesions were counted five days after inoculation. Five replicates were conducted for each sample.

For the curative assay *in vivo*[24,25], the whole leaves of *N. tabacum* L. cv. Samsun *NN* were inoculated with TMV particles (10.19 μg/mL final concentration). α and β-CBD were smeared onto the left half leaves 24 h after TMV inoculation, while the DMSO solution was smeared onto the right side as a negative control. The numbers of local lesions were counted five days after inoculation. Three replicates were conducted for each sample.

The inhibition rate of viral infection was calculated according to the following formula:

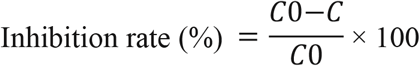

*C0*: the mean value of local lesions for the control treatment; *C*: the average number of local lesions for the treatments.

### 2.4. TMV-GFP infection foci measurement

The third leaf from the top of *N. benthamiana* plant was mechanically inoculated with TMV by rubbing the extracts of TMV-GFP infected leaves at 24 h after applying α(β)-CBD (75.0 μM) as foliar spray. Each treatment experiment was performed with five plants (replicates).

Fluorescent optical imaging of TMV-GFP inoculated *N. benthamiana* leaves was conducted under identical illumination and exposure conditions with an imaging fluorimeter Fluorcam FC 800-O (Photon System Instruments, Drasov, Czech Republic) at five and seven days post-inoculation (dpi), respectively. ImageJ v1.80 was used for the quantitative analysis of the GFP expression level and the areas of fluorescent infection sites [26].

### 2.5. Quantitative RT-PCR and TAS-ELISA

Total RNA was extracted with the MagMAX-96 Total RNA Isolation Kit (Thermo Fisher Scientific, Shanghai, China) following the user guide. cDNA was prepared by using the PrimeScript RT-PCR Kit (Takara Bio, Shiga, Japan). Quantitative RT-PCR (qRT-PCR) reactions were implemented with Platinum™ SYBR™ Green qPCR SuperMix - UDG (Invitrogen, Thermo Fisher Scientific) using the ABI 7500 real-time PCR system (Applied Biosystems, Carlsbad, CA). The reference gene *EF-1a* was used for quantitative analysis. The relative expression levels of the tested genes were calculated with the 2^−ΔΔCt^ method[27]. The primer sequences used are shown in Supplementary Table 1.

According to the previous studies[28], the accumulation of TMV protein was quantified by Triple antibody sandwich enzyme-linked immunosorbent (TAS-ELISA) assay kit (Agdia, Elkhart, IN, USA) following the user guide. was conducted to quantify in TMV infected tobacco leaves using an assay following the manufacturer’s instructions.

The inhibition level of viral proliferation was recorded and calculated according to the following formula:

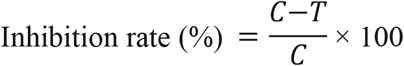

*C*: the viral content in the positive control; *T*: the viral content in the treated leaves.

Only TMV infected leaves were adopted as positive control. TMV concentration was calculated by the standard curve with the A_405_ value of TMV at concentrations of 8, 4, 2, 1, 0.5, 0.25 and 0.125 μg/mL. Absorbance at 405 nm was monitored with an iMark microplate reader (iMark13083, Bio-Rad, USA). All experiments were repeated three times and consisted of at least five tobacco seedlings per replicate.

### 2.6. Determination of JA and SA

The quantification of SA and JA from plant crude extracts were performed by LC-MS/MS following the previous description[29]. Approximately 200 mg harvested leaves were ground to a fine powder in liquid nitrogen. D6-SA (2-hydroxybenzoic acid-[2H6]) obtained from Sigma-Aldrich and H2-JA (dihydrojasmonic acid) obtained from OlChemim were used as internal standards. The supernatants after the removal of residues were analyzed on HPLC-tandem mass spectrometry (1200 L LC-MS system, Varian, American). Each treatment was consist of five replicates per sample.

### 2.7. Statistical analysis

All the experiment data presented with mean±SD were pooled across three independent repeated experiments. Analysis of variance (ANOVA) was performed using SPSS v 19.0 (SPSS Inc., Chicago, IL, USA). The statistical differences between α-CBD and β-CBD treatment were considered significant with a Student’s *t*-test at *P*<0.05.

## 3. Results

### 3.1. structural identification of CBD

To obtain α(β)-CBD with the high purity, an efficient and green extraction and purification method was established. In contrast to previous research, the extracts were first washed in 5% phosphate, rather than the mixed n-hexane/(methanol-water) solution, which has a remarkable effect on eliminating significant impurities including nicotine and water-soluble sugar. The prepared samples were then applied to the silicon column, and eluted by different ratios (1:1, 3:1, 5:1 and 7:1) of petroleum ether/ethyl acetate to determine which ratio was best for purifying CBDs and removing other components. The fractions eluted by a 1:1 ratio of petroleum ether/ethyl acetate had some CBTs, with a higher abundance around 50 min. However, there were still many other components (Fig. 2A). When the ratio was changed to 3:1, many CBTs were found, and the other components were decreased in quantity and abundance (Fig. 2B). The fractions eluted by a 3:1 ratio of petroleum ether/ethyl acetate had the largest number of CBDs than the other ratios (5:1 and 7:1), while the slightest number of other components was obtained in quantity and abundance (Fig. 2 C-D).

**Figure 2.**
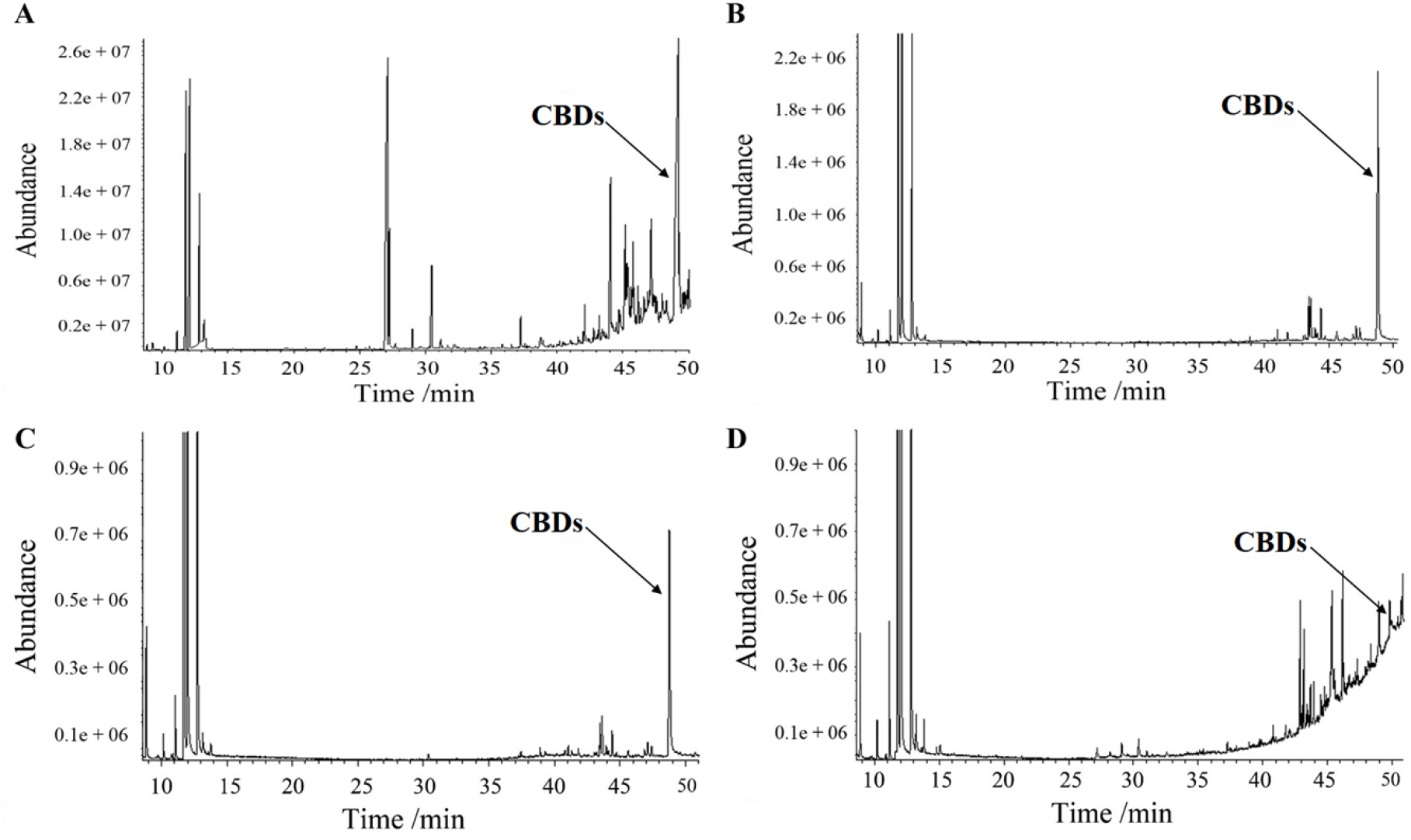
(A) Total ion current of fractions eluted by 1:1 ratio of petroleum ether/ethyl acetate. (B) Total ion current of fractions eluted by 3:1 ratio of petroleum ether/ethyl acetate. (C) Total ion current of fractions eluted by 5:1 ratio of petroleum ether/ethyl acetate. (D) Total ion current of fractions eluted by 7:1 ratio of petroleum ether/ethyl acetate.

We performed semi-preparative HPLC using the UPLC condition to obtain CBD with a 60:40 ratio of acetonitrile: water. Two distinct peaks emerged clearly within 90 sec. After the samples were eluted using the silicon column and applied to semi-preparative HPLC, two targeted peaks were entirely separated from other components and even the CBDs themselves within 35 min (Fig. 3–4). The purity of the two compounds done by semi-preparative HPLC was >98%, confirmed by UPLC (data not shown). The yield of compound 1 and compound 2 were 0.005% and 0.007%, respectively. The structures of the compounds 1 and 2 were shown in Fig. 1, and the ^1^H and ^13^C NMR data of 1 and 2 were listed in Figures S1–S4, respectively.

**Figure 3.**
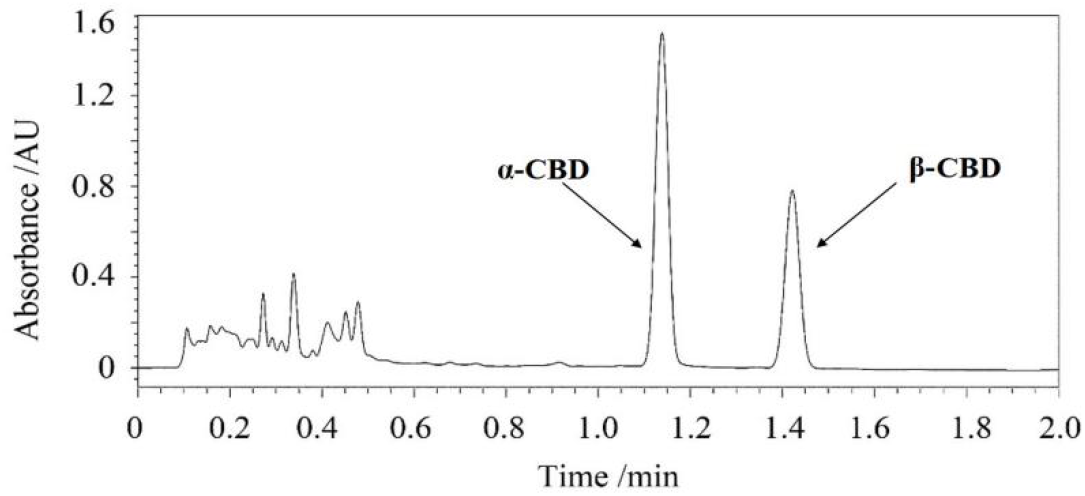
UPLC of mobile phase acetonitrile: water = 6:4 (v/v)

**Figure 4.**
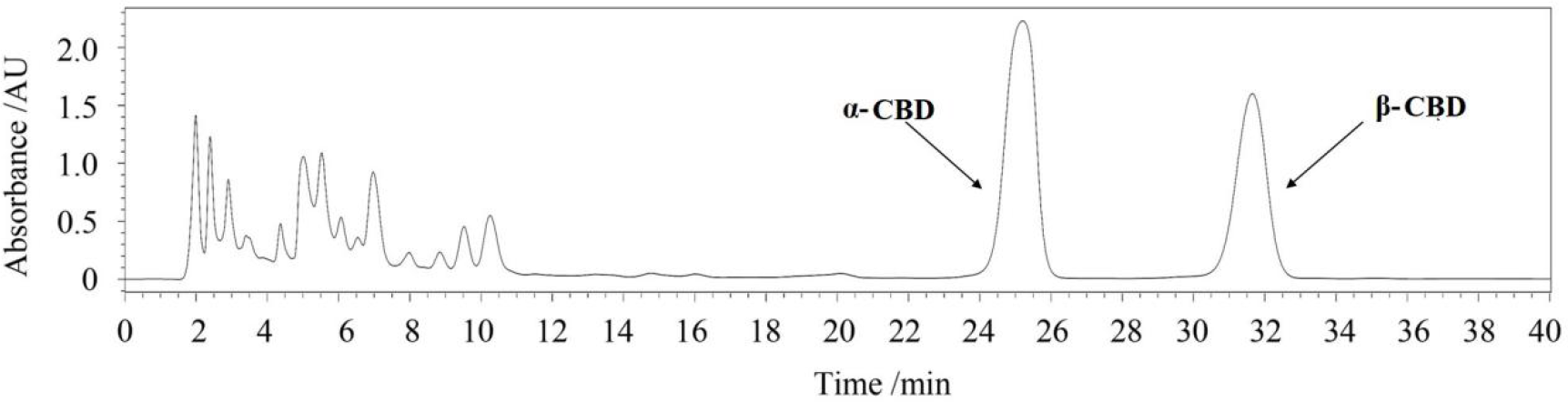
Semi-preparative HPLC of CBDs purified by C_18_ column.

The crystals of Peak 1, which melted at 65–66°C, were colorless. The crystals of Peak 2 were white, and the melting point was 125–126°C. The chemical structures of the two compounds were confirmed by high-resolution LC-MS and NMR, respectively. This experimental process is simple, easy to operate, and the obtained compounds have high purity, both above 98%.

The molecular formula of compound 1, C_20_H_34_O_2_, was deduced from MS, and its molecular weight was 306. Some useful information about compound 1 was established, including: m/z 329.24493 was [M+Na]^+^, m/z 289.25262 was [M-H_2_O+H]^+^ and m/z 635.50128 was [2M+Na]^+^ (Fig. 5A). In secondary MS of compound 1 (Fig. 5B), m/z 329.24 became 311.17 after losing one H_2_O. In triple MS, after losing both H_2_O and –C_3_H_6_, m/z 293.00 became 251.33 (Fig. 5C). Compound 1 was preliminarily determined as 2,7,11-cembratriene-4,6-diol. Both the fine structure and the absolute configuration of the chiral carbon atom at C_4_ of α-2,7,11-cembratriene-4,6-diol was confirmed by ^13^C-NMR and ^1^H-NMR (Figure S1-S2).

**Figure 5.**
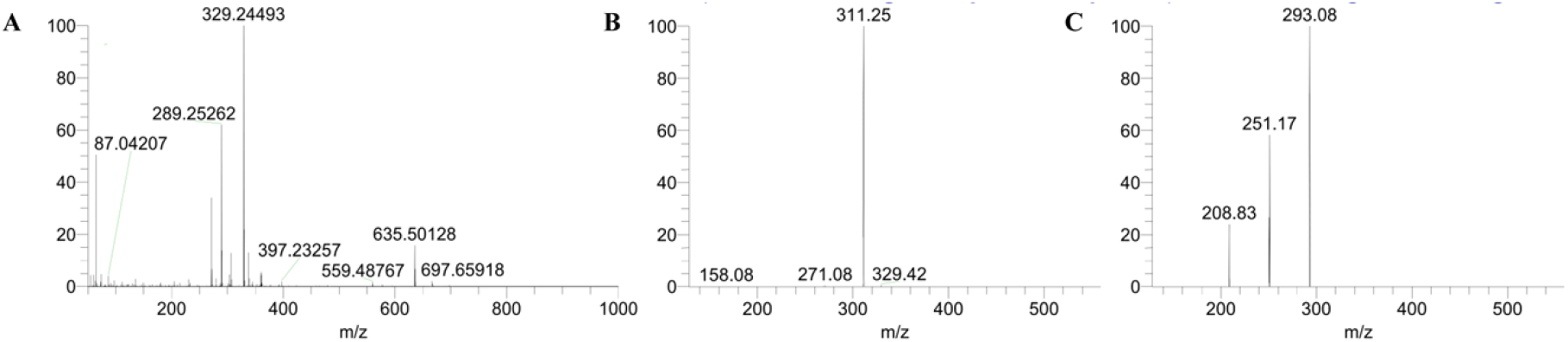
(A) High-resolution MS of Compound 1; (B) Secondary MS of Compound 1; (C) triple MS of Compound 1.

The molecular formula of compound 2, C_20_H_34_O_2_, was deduced from MS, and the molecular weight was 306. Some useful information about Compound 2 was established, including: m/z 329.24316 was [M+Na]^+^, m/z 289.25107 was [M-H_2_O+H]^+^ and m/z 635.49762 was [2M+Na]^+^ (Fig. 6A). In secondary MS of Compound 2 (Fig. 6B), m/z 329.24 became m/z 311.17 after losing one H_2_O. In triple MS, after losing both H_2_O and –C_3_H_6_, m/z 293.00 became 251.33 (Fig. 6C), which was similar to that of α-2,7,11-cembratriene-4,6-diol. Compound 2 was preliminarily determined as β-2,7,11-cembratriene-4,6-diol, and confirmed by ^13^C-NMR and ^1^H-NMR (Figure S3-S4).

**Figure 6.**
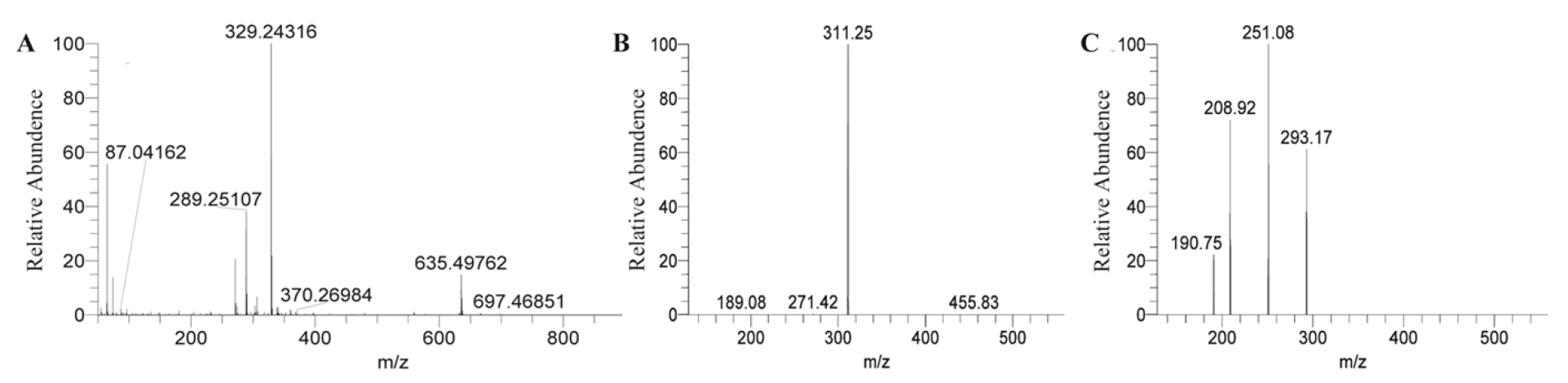
(A) High-resolution MS of Compound 2; (B) Secondary MS of Compound 2; (C) triple MS of Compound 1.

### 3.2. Exogenous CBDs Enhance Resistance to TMV

The local lesion host *N. tabacum* L. cv. Samsun *NN* pretreated with α- and β-CBD had fewer lesions than DMSO pretreated plants. As shown in Table 1, obvious inhibitory effects of α-CBD and β-CBD against TMV in a dose-dependent manner was obtained by the half-leaf method in *N. tabacum* L. cv. Samsun *NN*. α- and β-CBD treatments showed higher protective effects (73.2% and 71.6%) at 75.0 μg/mL than Ningnanmycin treatment (53.1%) at 200 μg/mL. Compared with the protective activity, α- and β-CBD at 75.0 μg/mL possessed relatively lower curative (35.0% and 38.7%) and inactive activities (37.4% and 33.2%).

**Table 1.** Inhibition effects of α and β-CBD on TMV in *N. tabacum* L. cv. Samsun *NN* by half leaf method. All results are expressed as means ± SD; Numbers with different letters are statistically different at *p* < 0.05, n = 3 for all groups.

| Treatment | Concentration ($\mu\text{g/mL}$) | Protection effect (%) | Curative effect (%) | Inactivation effect (%) |
| --- | --- | --- | --- | --- |
| α-CBD | 12.5 | 30.5± 7.6a | 21.4 ± 3a | 21.5 ± 4a |
| | 25.0 | 43.4 ± 4.9b | 28.1 ± 9b | 32.8 ± 6b |
| | 50.0 | 58.0 ± 5.3c | 33.0 ± 8c | 33.7 ± 5b |
| | 75.0 | 73.2 ± 4.2d | 35.0 ± 8c | 37.4 ± 4b |
| β-CBD | 12.5 | 38.0 ± 6.3b | 27.2 ± 6b | 25.3 ± 6a |
| | 25.0 | 45.9 ± 5.4b | 33.4 ± 3c | 27.0 ± 9ab |
| | 50.0 | 60.6 ± 7.1c | 36.5 ± 2cd | 30.5 ± 7b |
| | 75.0 | 71.6 ± 8.4d | 38.7 ± 8cd | 33.2 ± 4b |
| Ningnanmycin | 200 | 53.2 ± 5.5c | 51.4 ± 4d | 56.0 ± 6d |

TMV-GFP was used to analyze which virus infection steps involved in the inhibitory effects of α-CBD and β-CBD (Fig. 7). Compared with DMSO and Ningnanmycin treatments along with a higher GFP intensity, a significantly weaker viral GFP fluorescence signal and smaller infection area were obtained in α-CBD and β-CBD pretreated *N. benthamiana* (Fig. 7A). There are no clear differences in the foci number of TMV-GFP each plant and foci size per foci between α-CBD and β-CBD pretreatment (Fig. 7B and C). The meantime for TMV-GFP moving to apical leaves in α-CBD and β-CBD pretreatment was 2 days later than the Ningnanmycin treatment, although this difference was not statistically significant (*P*=0.08) (Fig. 7D).

**Figure 7.**
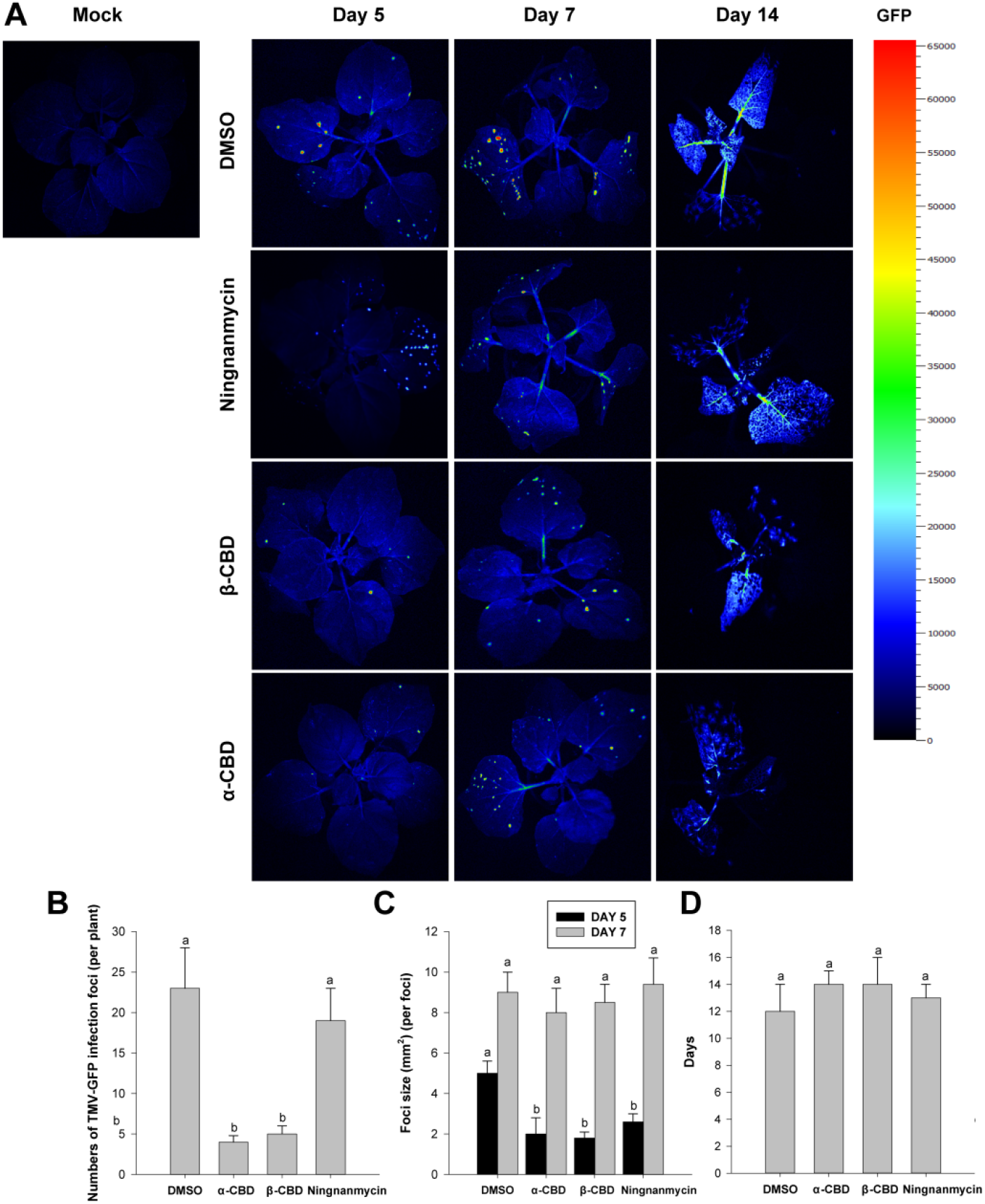
CBDs enhance N. benthamiana’ resistance against TMV. DMSO, Ningnanmycin, α-CBD and β-CBD pretreated plants were inoculated on the third leaf from top with TMV-GFP; (A) Fluorescent optical imaging of systemic TMV-GFP multiplication in different treatments was taken under UV light at 5, 7 and 14 days post-inoculation (dpi); (B) Numbers of infection foci on inoculated leaves; (C) TMV-GFP multiplication was quantified in the treated plants by calculating the foci size per foci; (D) The mean time for TMV-GFP moving to apical leaves in each treatment. Statistical significance was tested by the Kruskal–Wallis test and significantly different groups are indicated by different letters for each time point (P< 0.05). The data were the mean ± SD of three replicates (at least 20 plants per replicate). Similar results were obtained in three independent repeats of the experiment.

The results of qRT-PCR indicated that any distinct differences in the relative expression of the TMV-CP gene were not detected among the control and both CBDs treatments, while an approximately 55% decrease of the TMV-CP gene in Ningnanmycin treatment was obtained (Fig. 8A). However, TAS-ELISA results showed that the viral protein synthesis was prominently inhibited in CBDs treated leaves (Fig. 8B). At 75 μg/mL, α-CBD and β-CBD treatments caused 50% and 48% decreases in the contents of TMV-CP protein compared to the control, respectively, while 22% reduce in the viral protein content was gained in Ningnanmycin treatment (Fig. 8B and Table 2). These results suggest that CBDs may mainly obstruct the virus protein biosynthesis processes.

**Figure 8.**
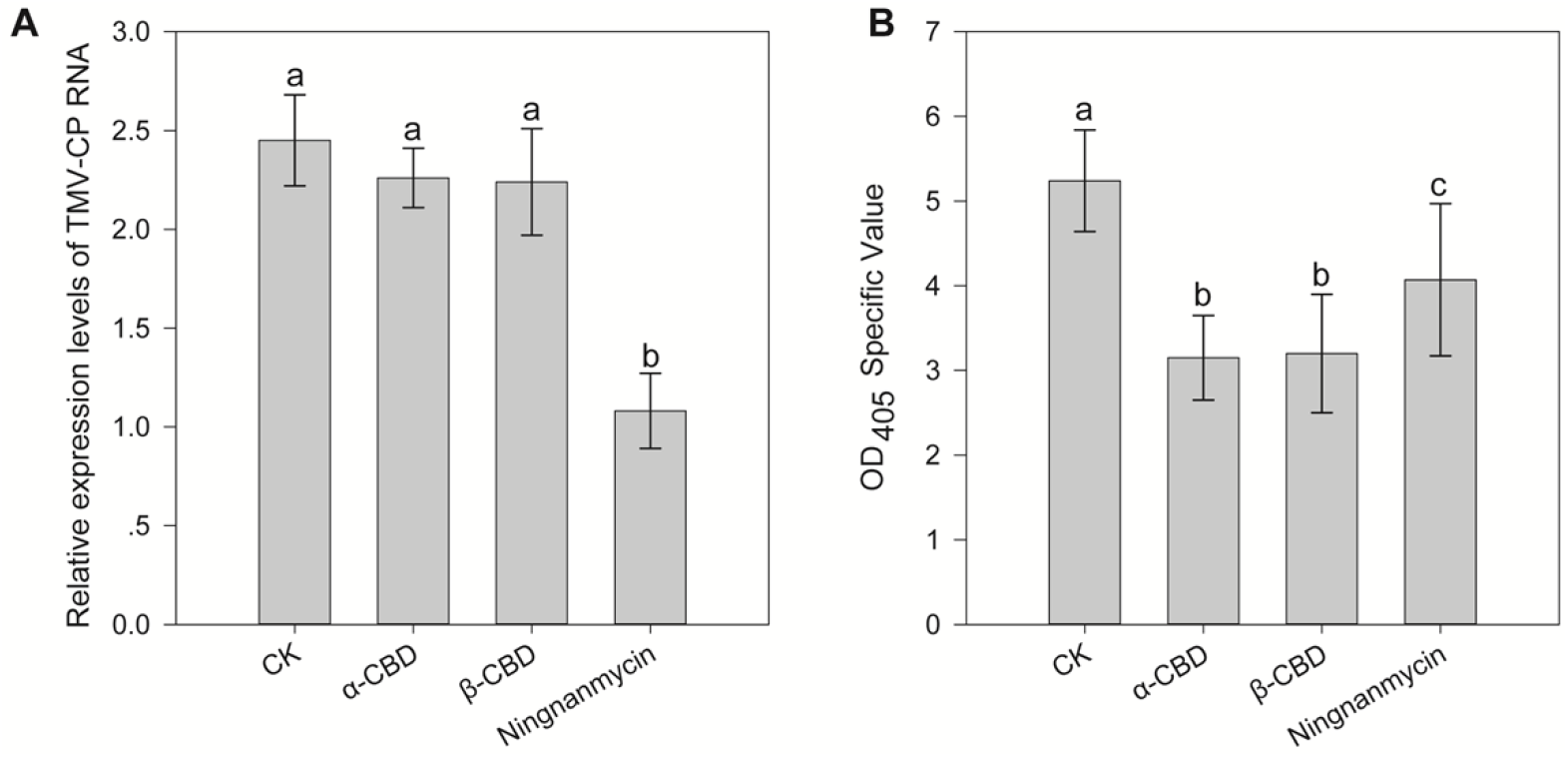
Protective effects of α and β-CBD in N. benthamiana on TMV. (A) Gene expression Levels of TMV CP in α and β-CBD-treated leaves; (B) Relative accumulation of TMV protein in tobacco leaves in α and β-CBD-treated leaves.

**Table 2.** Inhibition effects of α and β-CBD in N. benthamiana on TMV by TAS-ELISA. All results are expressed as means ± SD; Numbers with different letters are statistically different at p < 0.05, n = 3 for all groups.

| Sample | Concentration ($\mu\text{g/mL}$) | Protective effect (%) | Inactive effect (%) | Curative effect (%) |
| --- | --- | --- | --- | --- |
| α-CBD | 75 | 68.7 ± 3a | 36.4± 5a | 34.1± 4a |
| β-CBD | 75 | 65.2 ± 5a | 39.2± 3a | 32.2± 3a |
| Ningnanmycin | 200 | 51.3 ± 2b | 56.7± 5b | 47.5± 3a |

### 3.3. Effects of CBDs on phytohormone levels

The concentrations of JA in the CBDs-treated leaves under TMV infection were almost 1.8 times at 24 hpi and 1.5 times at 36 hpi higher than that of the control treatments, respectively, while the contents of SA in the treated leaves were almost 2.0 times at 24 hpi and 1.4 times at 36 hpi higher (Fig. 9). Meanwhile, no significant differences were obtained between α-CBD and β-CBD treatments. These results implied that CBDs could evidently enhance the generation of JA and SA in the treatment leaves.

**Figure 9.**
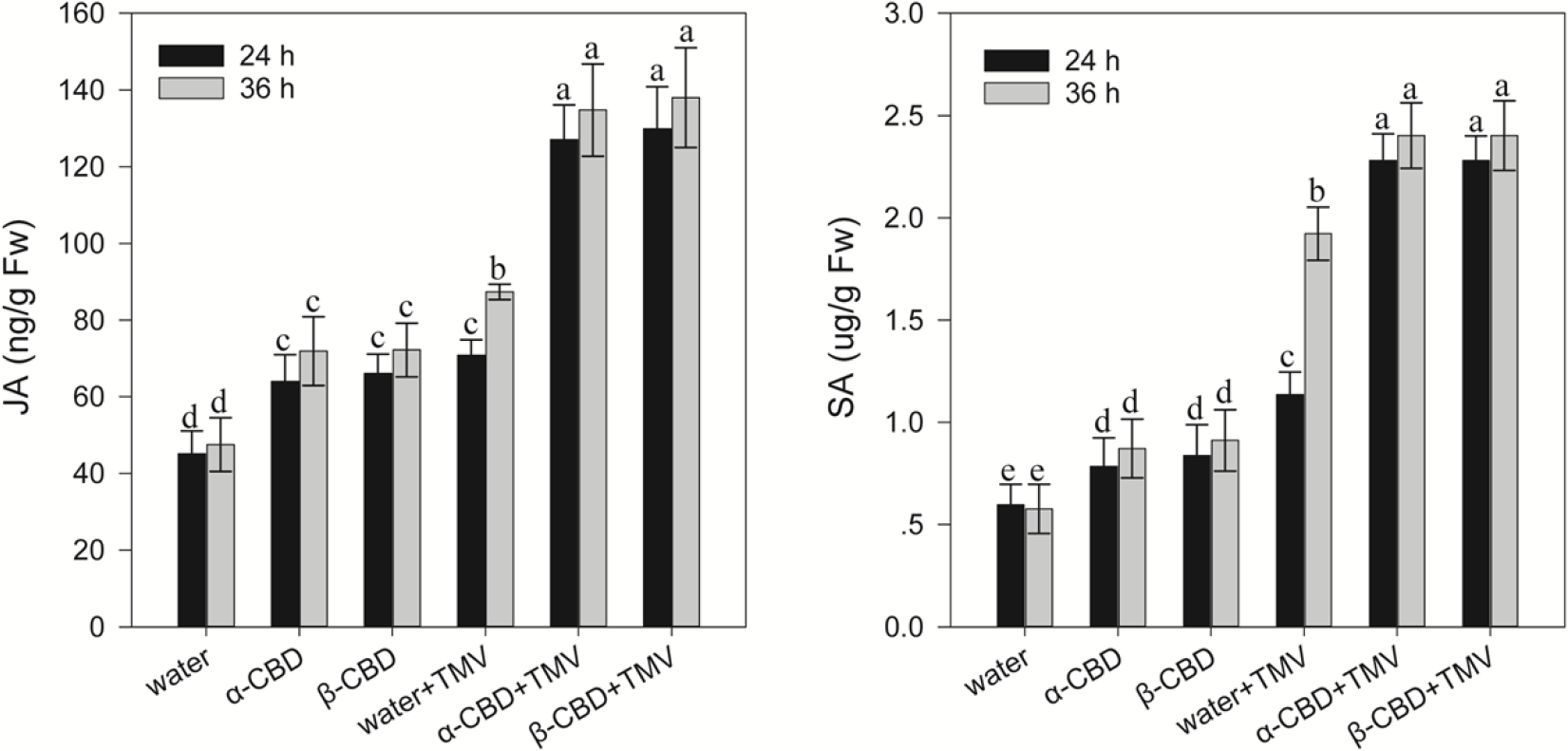
Effect of α-CBD and β-CBD on the contents of JA and SA at 24 h and 36 h after CBD treatment under TMV infection.

**Figure 10.**
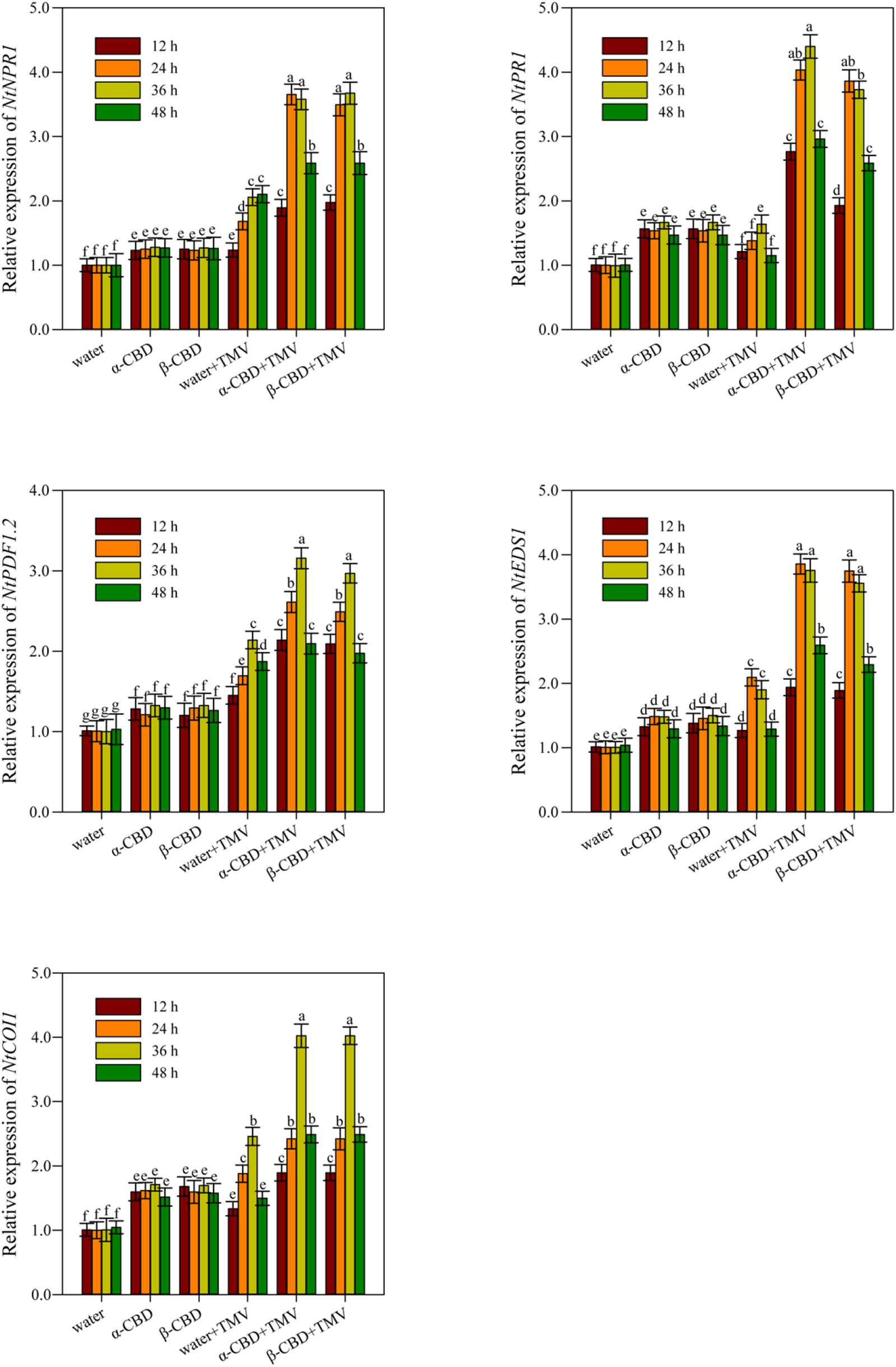
Relative expression of key genes of JA (COI1 and PDF1.2) and SA signaling pathway genes (PR1, NPR1 and EDS1).

The transcription of JA signaling pathway (*COI1* and *PDF1.2*) and SA signaling pathway genes (*PR1*, *NPR1* and *EDS1*) has made for obvious differences in performance. The larger elevation and longer duration of all these marker genes expression in CBDs pretreated plants occurred than that in the water pretreatment under TMV stress. The highest expression levels of JA and SA signaling pathway genes were observed at 24 hpi and 36 hpi. However, there were no significant differences in the expression of the tested genes between α-CBD+ TMV and β-CBD+TMV treatments. These transcriptions have similar trends in the diversification of JA and SA (Fig. 3G and H).

## 4. Discussion

Cembranoids are carbocyclic diterpenes and have gained widespread attention for the past few years and become one of the main focuses on natural product chemistry[11]. In this study, we successfully obtained α(β)-CBD with high purity (98%) using an efficient and green extraction and purification method. Compared with previous reports, the present method eliminates liquid-liquid extraction process of the n-hexane/(methanol-water) mixed solution, and innovatively added 5% phosphoric acid to wash for 1 to 2 times, then washed with deionized water for 2 to 3 times to neutral in impurities process, which has a significant effect on removing major impurities such as nicotine and water-soluble sugars. Furthermore, the present method is simple and easy to operate. Large amounts of high-purity compounds can be obtained after one time silica gel chromatographic column separation and impurity removal and further purification by HPLC method.

Several natural products extracted from *N. tabacum* exhibited anti-TMV activity, such as flavonoids from the roots and stems of Yunyan-202[6], biphenyls from the leaves of *Honghuadajinyuan*[7], sesquiterpenes and benzolactones from the leaves of Yunyan-201[8]. Here, we found that α(β)-CBD from Yunyan 100 leaves exhibited higher protection effects than the curative and inactive activities in *N. tabacum* L. cv. Samsun *NN* by the half-leaf process in compared with the previously mentioned anti-TMV components from different varieties of tobacco. For initial screening, TMV and its local lesion host *N. tabacum* L. cv. Samsun *NN* was used. However, sometimes only the counts of local lesion for the screen of antiviral agents may not be enough in susceptible plants. Therefore, *N. benthamiana* for tracking assay by green fluorescent-protein tagged TMV. The tendency of their antiviral activities was confirmed by ID-ELISA in K326 and tracking images of TMV-GFP in *N. benthamiana*.

CP is essential for viral systemic infection and replication, which is involved in cell to cell and long-distance transport of TMV and the host response[30]. Our qRT-PCR and TAS-ELISA results indicated that α(β)-CBD significantly interfere with either TMV CP protein biosynthesis or post-translational modification processes, which is consistent with the observed TMV-GFP images of inoculated *N. benthamiana* leaves. However, full exploration are still needed whether CBD directly affect the structure of virus particle and stereoscopic assembly of the virus.

Besides, α(β)-CBD can elicit plant defense against virus infection without cost to the plant’s fitness. Much of previous researches have reported that tobacco diterpenes play divers roles, functioning as signaling molecules, or activating agents in plant defense. Seo et al. (2003) documented that Z-abienol-related diterpene (11E,13E)-labda-11,13-diene-8α,15-diol could stimulate tobacco plants defense response against TMV infection[5]. Fujimoto et al. (2015) showed that sclareol as an elicitor-related substance boosts the resistance of *Arabidopsis* and tomato roots against nematode penetration[31]. The signaling molecules SA and JA play critical roles in the defense response of tobacco plants to TMV. The enhanced production of SA and JA and the significantly increased transcription of the tested genes *COI1, PDF1.2, PR1*, *NPR1* and *EDS1* further conformed that exogenously applied α(β)-CBD can effectively elicit the tobacco plant immunity against TMV.

In conclusion, α-CBD and β-CBD with high purity (98%) were separated and purified from tobacco leaves using an efficient and green extraction and purification method. Exogenously applied α(β)-CBD showed excellent inhibitory activity against TMV infection in a dose-dependent manner. In particular, α- and β-CBD at 75.0 μg/mL exhibited higher protective effects with the values of 73.2% and 71.6% than that of Ningnanmycin (53.1%) at 200 μg/mL. The significant antiviral activity is due to its ability to disturb either TMV CP protein biosynthesis or post-translational modification processes and active natural immunity of tobacco plants by inducing the increased activity of defense-related enzymes and the improved transcription of SA signaling pathway genes. Collectively, α(β)-CBD have potential as new disease resistance elicitors to control TMV in the future.

## Author Contributions

Y.-X.F., H.-J.J. and H.G. isolated and purified of active compounds and analyzed the structure. W.-D. Z., Y.-B. Z., M. A. S. tested anti-TMV activities of α-CBD and β-CBD. D.-S.G. tested Transmission Electron Microscope (TEM). W.D. Z., Y.B. Z., K.Y. L., Y.J. W., Q.L. L., W.H. C. determined the defense-related enzymes activity and the relative expression of SA and JA signaling pathway genes. S.-H. and J.-Y.W. designed the experiments and supervised the research. J. W., J.-Y.W. and Y.-X.F. wrote the manuscript. All authors have read and agreed to the published version of the manuscript.

## Supporting information materials

**Table S1. Groups of real-time quantitative PCR (RT-qPCR) primers used to amplify genespecific regions.**

**Figure S1. 13C-NMR spectra of α -CBD**

**Figure S2. 1H-NMR spectra of α -CBD**

**Figure S3. 13C-NMR spectra of β -CBD**

**Figure S4. 1H-NMR spectra of β -CBD**

## Reference

1. Jassbi, A. R.; Zare, S.; Asadollahi, M.; Schuman, M. C. Ecological Roles and Biological Activities of Specialized Metabolites from the Genus Nicotiana. Chem. Rev. 2017, 117 (19), 12227–12280.

2. Uzelac, B.; Stojičić, D.; Budimir, S. Glandular Trichomes on the Leaves of *Nicotiana tabacum*: Morphology, Developmental Ultrastructure, and Secondary Metabolites. In Plant Cell and Tissue Differentiation and Secondary Metabolites: Fundamentals and Applications, Ramawat, K. G.; Ekiert, H. M.; Goyal, S., Eds. Springer International Publishing: Cham, 2021; pp 25–61.

3. Steede, W. T.; Ma, J. M.; Eickholt, D. P.; Drake-Stowe, K. E.; Kernodle, S. P.; Shew, H. D.; Danehower, D. A.; Lewis, R. S. The tobacco trichome exudate z-abienol and its relationship with plant resistance to *phytophthora nicotianae*. Plant Dis. 2017, 101 (7), 1214–1221.

4. Scalabrin, E.; Radaelli, M.; Rizzato, G.; Bogani, P.; Buiatti, M.; Gambaro, A.; Capodaglio, G. Metabolomic analysis of wild and transgenic *Nicotiana langsdorffii* plants exposed to abiotic stresses: unraveling metabolic responses. Anal. Bioanal. Chem. 2015, 407 (21), 6357–6368.

5. Seo, S.; Seto, H.; Koshino, H.; Yoshida, S.; Ohashi, Y. A diterpene as an endogenous signal for the activation of defense responses to infection with tobacco mosaic virus and wounding in tobacco. Plant cell 2003, 15, 863–73.

6. Wang, X.; Huang, L. J.; Liang, M. J.; Li, Y. K.; Zeng, W. L.; Xiang, H. Y.; Li, J.; Liu, X.; Mi, Q. L.; Guo, Y. D.; Yang, G. Y.; Deng, L.; Gao, Q. Two new furan-2-carboxylic derivatives from the leaves of *Nicotiana tabacum* and their anti-tobacco mosaic virus activities. Chem. Nat. Compd. 2020, 56 (5), 848–851.

7. Islam, W.; Qasim, M.; Noman, A.; Tayyab, M.; Chen, S.; Wang, L. Management of tobacco mosaic virus through natural metabolites. Rec. Nat. Prod. 2018, 12 (5), 403–415.

8. Shang, S. Z.; Duan, Y. X.; Zhang, X.; Pu, J. X.; Sun, H. D.; Chen, Z. Y.; Miao, M. M.; Yang, G. Y.; Chen, Y. K. Phenolic amides from the leaves of *Nicotiana tabacum* and their anti-tobacco mosaic virus activities. Phytochem. Lett. 2014, 9, 184−187.

9. El Sayed, K.A., Sylvester, P.W. Biocatalytic and semisynthetic studies of the anticancer tobacco cembranoids. Expert Opin. Invest. Drugs 2007, 16, 877–887.

10. Liu, X.; Zhang, J.; Liu, Q.; Tang, G.; Wang, H.; Fan, C.; Yin, S. Bioactive cembranoids from the south China sea soft coral *Sarcophyton elegans*. Molecules 2015, 20, 13324–13335.

11. Yan, N.; Du, Y.; Liu, X.; Zhang, H.; Liu, Y.; Zhang, P.; Gong, D. P.; Zhang Z. F. Chemical structures, biosynthesis, bioactivities, biocatalysis and semisynthesis of tobacco cembranoids: An overview. Ind. Crops Prod. 2016, 83, 66–80

12. Aqil, F.; Zahin, M.; El Sayed, K. A.; Ahmad, I.; Orabi, K. Y.; Arif, J. M. Antimicrobial, antioxidant, and antimutagenic activities of selected marine natural products and tobacco cembranoids. Drug Chem. Toxicol. 2011, 34, 167–179.

13. Kennedy, B. S.; Nielsen, M. T.; Severson, R. F. Biorationals from Nicotiana protect cucumbers against *Colletotrichum lagenarium* (Pass.) ell. & halst disease development. J. Chem. Ecol. 1995, 21, 221–231.

14. Kennedy, B. S.; Nielsen, M. T.; Severson, R. F.; Sisson, V. A.; Stephenson, M. K.; Jackson, D. M. Leaf surface chemicals from Nicotiana affecting germination of *Peronospora tabacina* (adam) sporangia. J. Chem. Ecol. 1992, 18, 1467–1479.

15. Duan, S.; Du, Y.; Hou, X.; Li, D.; Ren, X.; Dong, W.; Zhao, W.; Zhang, Z. Inhibitory effects of tobacco extracts on eleven phytopathogenic fungi. Nat. Prod. Res. Dev. 2015, 27, 470–474.

16. Yan, N.; Du, Y.; Liu, X.; Zhang, H.; Liu, Y.; Shi, J.; Xue, S.J.; Zhang, Z. Analyses of effects of α-cembratrien-diol on cell morphology and transcriptome of *Valsa mali* var. mali. Food Chem. 2017, 214, 110–118.

17. Sui, J.; Wang, C.; Liu, X.;Fang, N.; Liu, Y.; Wang, W.; Yan, N.; Zhang, H.B.; Du, Y.; Liu, X.; Lu, T.; Zhang, Z.; Zhang, H. Formation of alpha- and beta-Cembratriene-Diols in tobacco (*nicotiana tabacum* l.) is regulated by jasmonate-signaling components via manipulating multiple cembranoid synthetic genes. Molecules 2018, 23, 10

18. Ren, X.; He, X.; Zhang, Z.; Yan, Z.; Jin, H.; Li, X.; Qin, B. Isolation, identification, and autotoxicity effect of allelochemicals from rhizosphere soils of flue-cured tobacco. J. Agric. Food. Chem. 2015, 63, 8975–8980.

19. He, X.; Hou, X.; Ren, X.; Guo, K.; Li, X.; Yan, Z.; Du, Y.; Zhang, Z.; Qin, B. Two new cembranic diterpenoids from the flowers of *Nicotiana tabacum* L. Phytochem. Lett. 2016, 15, 238–244.

20. Marshall, J. A.; Robinson, E. D. Enantioselective total synthesis of (+)-α-2, 7, 11-cembratriene-4, 6-diol(α-CBT). Tetrahedron Lett. 1989, 30(9):1055–1058.

21. Marshall, J. A.; Robinson, E. D.; Lebreton, J. Synthesis of the tumor-inhibitory tobacco constituents α and β-2,7,11-cembratriene-4,6-diol by diastereoselective [2,3] Wittig ring contraction. J. Org. Chem. 1990, 55(1), 227–239.

22. Gooding, G. V.; Hebert, T. T. A simple technique for purification of tobacco mosaic virus in large quantities. Phytopathology 1967, 57, 1285.

23. Li, L.; Zou, J.; Xu, C.; You, S.; Li, Y.; Wang, Q. Synthesis and Anti-Tobacco Mosaic Virus/Fungicidal/Insecticidal/Antitumor Bioactivities of Natural Product Hemigossypol and Its Derivatives. J. Agric. Food Chem. 2021, 69 (4), 1224–1233.

24. Wang, J.; Yu, G.; Li, Y.; Shen, L.; Qian, Y.; Yang, J.; Wang, F. Inhibitory effects of sulfated lentinan with different degree of sulfation against tobacco mosaic virus (TMV) in tobacco seedlings. Pestic. Biochem. Phys. 2015, 122, 38–43.

25. Abdelkefi, H.; Sugliani, M.; Ke, H.; Harchouni, S.; Soubigou-Taconnat, L.; Citerne, S.; Mouille, G.; Fakhfakh, H.; Robaglia, C.; Field, B. Guanosine tetraphosphate modulates salicylic acid signalling and the resistance of *Arabidopsis thaliana* to Turnip mosaic virus. Mol. Plant Pathol. 2018, 19 (3), 634–646.

26. Cai, L.; Zhang, W.; Jia, H.; Feng, H.; Wei, X.; Chen, H.; Wang, D.; Xue, Y.; Sun, X. Plant-derived compounds: A potential source of drugs against Tobacco mosaic virus. Pestic. Biochem. Phys. 2020, 169, 104589.

27. Liu, H.; Zhao, X.; Yu, M.; Meng, L.; Zhou, T.; Shan, Y.; Liu, X.; Xia, Z.; An, M.; Wu, Y. Transcriptomic and functional analyses indicate novel anti-viral mode of actions on tobacco mosaic virus of a microbial natural product ∊-Poly-l-lysine. J. Agric. Food Chem. 2021, 69 (7), 2076–2086.

28. Zhu, F.; Zhang, P.; Meng, Y. F.; Xu, F.; Zhang, D. W.; Cheng, J.; L, H.H.;X, D. H. Alpha-momorcharin, a RIP produced by bitter melon, enhances defense response in tobacco plants against diverse plant viruses and shows antifungal activity *in vitro*. Planta 2013, 237, 77–88.

29. Pan, X.; Welti, R.; Wang, X. Quantitative analysis of major plant hormones in crude plant extracts by high-performance liquid chromatography–mass spectrometry. Nat. Prot. 2010, 5, 986–992.

30. Asurmendi, S.; Berg, R.H.; Koo, J.C.; Beachy, R.N. Coat protein regulates formation of replication complexes during tobacco mosaic virus infection. Proc. Natl. Acad. Sci. USA 2004, 101, 1415–1420.

31. Fujimoto, T.; Mizukubo, T.; Abe, H.; Seo, S. Sclareol induces plant resistance to root-knot nematode partially through ethylene-dependent enhancement of lignin accumulation. Mol Plant Microbe Interact. 2015, 28(4), 398–407.

